# Hmx gene conservation identifies the evolutionary origin of vertebrate cranial ganglia

**DOI:** 10.1101/2020.09.07.281501

**Authors:** Vasileios Papdogiannis, Hugo J. Parker, Alessandro Pennati, Cedric Patthey, Marianne E. Bronner, Sebastian M. Shimeld

**Affiliations:** Department of Zoology, University of Oxford, 11a Mansfield Road, Oxford OX1 3SZ; Stowers Institute for Medical Research, Kansas City, MO 64110, USA; Department of Radiosciences, Umeå University, 901 85 Umeå, Sweden; Division of Biology and Biological Engineering, California Institute of Technology, Pasadena, CA 91125, USA

## Abstract

The evolutionary origin of vertebrates included innovations in sensory processing associated with the acquisition of a predatory lifestyle^1^. Vertebrates perceive external stimuli through sensory systems serviced by cranial sensory ganglia (CSG) which develop from cranial placodes; however understanding the evolutionary origin of placodes and CSGs is hampered by the gulf between living lineages and difficulty in assigning homology between cell types and structures. Here we use the Hmx gene family to address this question. We show Hmx is a constitutive component of vertebrate CSG development and that *Hmx* in the tunicate *Ciona* is able to drive the differentiation program of Bipolar Tail Neurons (BTNs), cells previously thought neural crest homologs^2,3^. Using *Ciona* and lamprey transgenesis we demonstrate that a unique, tandemly duplicated enhancer pair regulated Hmx in the stem-vertebrate lineage. Strikingly, we also show robust vertebrate Hmx enhancer function in *Ciona*, demonstrating that deep conservation of the upstream regulatory network spans the evolutionary origin of vertebrates. These experiments demonstrate regulatory and functional conservation between *Ciona* and vertebrate *Hmx*, and confirm BTNs as CSG homologs. Our analysis also identifies derived evolutionary changes, including a genetic basis for secondary simplicity in *Ciona* and unique regulatory complexity in vertebrates.

---

CSG, including the trigeminal, vestibuloacoustic and epibranchial ganglia, relay information from sensory cells to the brain. CSG neurons derive from two sources: cranial placodes provide neurons that delaminate from the cranial ectoderm, and cranial neural crest cells migrate into ganglia providing all of the glia and some neurons of the trigeminal ganglia. The evolution of neural crest, placodes and CSG form part of the influential ‘New Head Hypothesis’ which posits that these and other innovations underlie the transformation of an ancestral chordate filter feeder into the ancestral vertebrate-type predator^1^. Our molecular and genetic understanding of this transformation has been limited, however, by the substantial anatomical gulf between vertebrates and their nearest living relatives, amphioxus and tunicates, which lack most or all of these characters (Fig. 1A)^4^.

**Figure 1.**
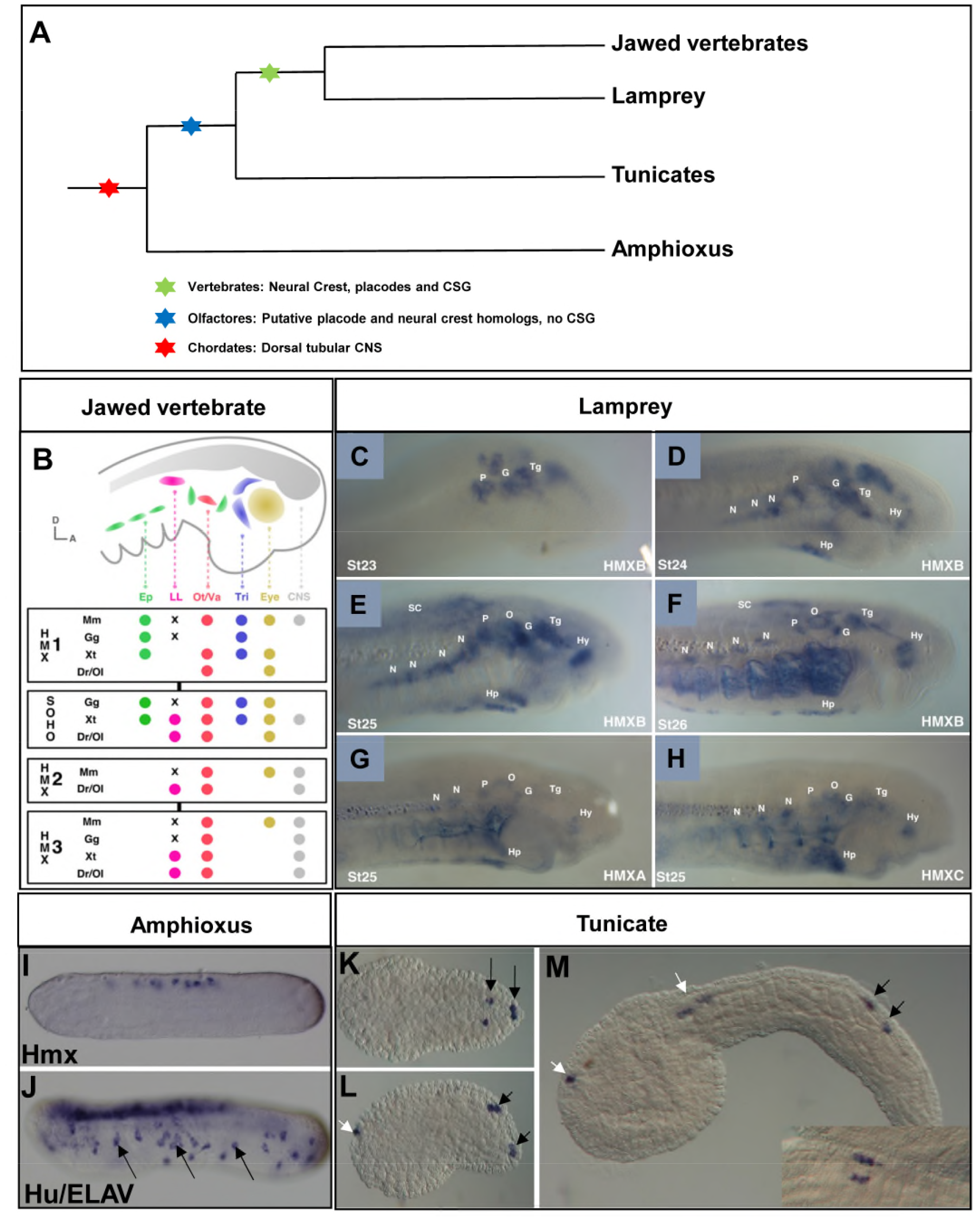
HMX expression in chordates. **A.** Phylogeny of the chordates showing the evolutionary ancestry of key characters. **B.** Schematic depiction of Hmx expression in jawed vertebrate cranial placodes/ganglia. Species shown are: Mm, *Mus musculus.* Gg, *Gallus gallus.* Xt-*Xenopus tropicalis.* Dr, *Danio rerio.* Ol, *Oryzias latipes.* Where species are missing it means either the gene has not been analysed (*Hmx2* in Gg and Xt) or the gene has been lost (*SOHo* in Mm). Other abbreviations: Ep, epibranchial. LL, lateral line. Ot/Va, otic/vestibuloacoustic. Tg, trigeminal. CNS, central nervous system. X indicates lateral line ganglia have been lost by these species. The olfactory is not shown as Hmx has not been reported from this placode. Full analysis underlying this figure in Supplementary Figure S4. **C-H** Expression of lamprey *Hmx* genes. **C.** *HmxB* expression starts in cranial sensory ganglia at stage 23. **D.** Hypothalamus expression in also seen by stage 24. An expression domain below the branchial basket corresponds to the position of the hypobranchial ganglia. **F.** Hindbrain and spinal cord are strongly marked after stage 25. **G, H**. Expression of lamprey *HmxA* and *HmxC* is identical to *HmxB.* Abbreviations: N, nodose. P, petrosal. G, geniculate. O, otic. Tg, trigeminal. SC, spinal cord. Hp, hypobranchial. Hy, hypothalamus **I, J.** *Hmx* expression in amphioxus compared to the neural marker *Hu/ELAV.* Arrows mark some of the peripheral neurons expressing *Hu/ELAV.* **K-M.** *Hmx* expression In *Ciona.* Black arrows identify BTNs, white arrows expression in the CNS. The inset on M shows a dorsal view with BTNs lying parallel to the CNS.

To address this gap we focused on the Hmx gene family, which encodes homeodomain transcription factors. We previously used transcriptomics to find markers for placode-derived CSG neurons, identifying *Hmx3* as one such gene^5^. Jawed vertebrates have 4 Hmx family genes named *Hmx1, Hmx2, Hmx3* and *SOHo*^6^, with expression in mouse, chicken, *Xenopus* and zebrafish primarily confined to the central nervous system (CNS), as well as cranial placodes and the CSG they form (Fig. 1B)^6–13^. Lampreys are members of the earliest-diverging living vertebrate lineage and offer insight into the basal vertebrate state. We identified three Hmx genes in lamprey, *HmxA, HmxB* and *HmxC*, all expressed in the CNS, cranial placodes and CSG (Fig. 1C-H). Lamprey Hmx genes were not expressed in the olfactory placode or in other parts of the peripheral nervous system (PNS) such as the dorsal root ganglia (DRG). This suggests that expression of vertebrate Hmx genes in the CNS, posterior placodes and their descendent CSG reflects the ancestral condition. To gain insight into roles of Hmx genes in invertebrate chordates, we next examined the expression of the single *Hmx* genes found in amphioxus^14^ and tunicates^15^. Amphioxus *Hmx* was expressed in the CNS but not in the PNS (Fig. 1I,J). In the tunicate *Ciona intestinalis, Hmx* was expressed in the CNS and in a subpopulation of PNS cells, the BTNs (Fig. 1K-M). BTNs are born lateral to the neural plate, and delaminate and migrate before connecting epidermal sensory cells to the CNS^3,16^. Previously, they have been likened to neural crest^2,3^.

Vertebrate Hmx genes are necessary for the correct development of CSG^11,17–19^. To test whether *Ciona Hmx* functions in BTN development, we used the *Ciona epiB* promoter^20^ to drive *Hmx* expression broadly in the embryonic epidermis, with transgenic embryos subsequently prepared for transcriptome analysis. Comparison of experimental to control embryos (Fig. 2A) revealed up- and down-regulated genes that were compared to cell-type expression profiles extracted from single-cell sequence data^2^. The results showed that *Ciona Hmx* upregulated the BTN differentiation program (Fig. 2B), concomitantly suppressing expression of epidermal genes, and of genes associated with other cells that develop from this lineage (Fig. 2B). This suggests that *Hmx* is sufficient to drive the BTN transcriptional program in *Ciona* embryonic epidermis.

**Figure 2.**
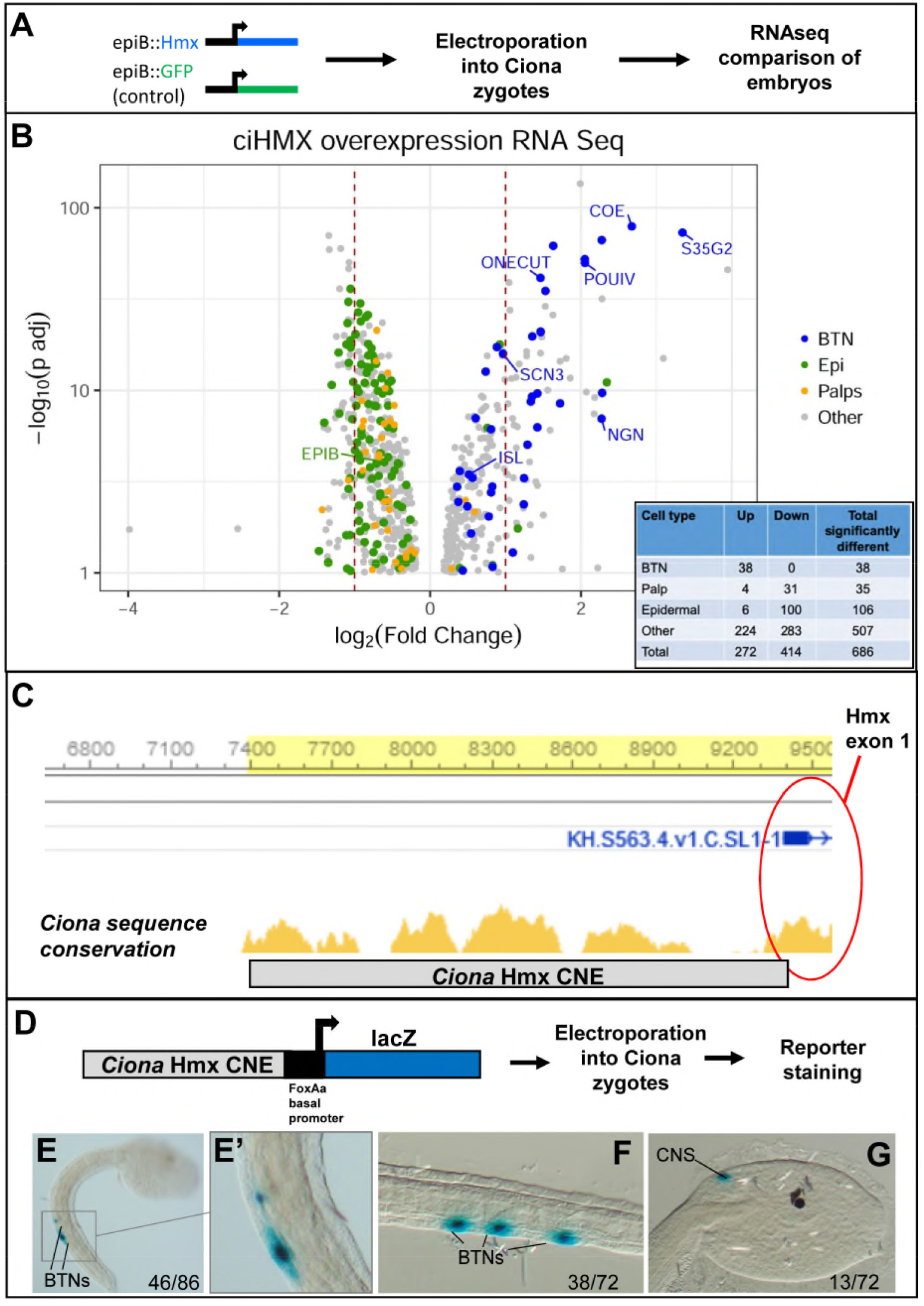
*Hmx* function and regulation in *Ciona*. **A.** Schematic of the experimental strategy used to drive *Ciona Hmx* overexpression. **B.** Volcano plot of genes up or down regulated after *Ciona Hmx* overexpression. Selected genes are named. Colour coding reflects genes identified as cell type expressed in single cell sequencing data^2^, and the inset table shows these quantified for BTN, palp and epidermal (EPI) cell types. **C.** Sequence conservation between *C. robusta* and *Ciona savignyi* identifies an approximately 2Kb CNE 5’ to the first *Hmx* exon. **D.** Schematic of the experimental strategy for characterising *Hmx* CNE activity in *Ciona.* **E-G.** *Ciona Hmx* CNE activity in *Ciona* embryos, visualised by lacZ staining. E shows a tailbud stage embryo with stain in BTNs, seen in close up in E’. No CNS stain was observed at this stage. F and G show BTN and CNS staining respectively in early larvae. Numbers indicate the number of embryos showing staining in the indicated structures, out of the total number of surviving embryos. Full embryo counts are in Supplementary Figure S5.

The expression domains and developmental roles of Hmx genes in vertebrates and *Ciona* may reflect either shared ancestry, or derived traits in either lineage. Discriminating between these possibilities requires comparison of the genetic program(s) upstream of Hmx expression in each lineage. To address this we mapped and tested Hmx regulatory elements in *Ciona* and lamprey. Comparison of tunicate genomes^21^ showed 2kb of sequence 5’ to the *Hmx* transcription start site to be conserved between *Ciona* species (Fig. 2C). We hypothesized this was a conserved non-coding element (CNE), with sequence conservation reflecting evolutionary constraint deriving from a role in gene regulation. We tested this in transgenic *Ciona* (Fig. 2D), and found the 2kb fragment able to drive robust and specific reporter expression in BTNs and in part of the CNS, recapitulating aspects of endogenous *Hmx* expression (Fig. 2E-G).

Hmx gene evolution in vertebrates is more complex. Jawed vertebrate Hmx genes are located in two paralagous two-gene clusters^6^. We found that the jawless vertebrate Hmx genes are in a single three-gene cluster in both lamprey and hagfish genomes (Fig. 3A, B). Sequence comparison shows these genomic arrangements evolved by gene duplication (Fig. 3E). In the vertebrate ancestor a single Hmx gene duplicated in tandem to yield a two-gene cluster. In jawless vertebrates a second tandem duplication occurred, yielding the three-gene cluster state found in lamprey and hagfish. In jawed vertebrates the two-gene cluster duplicated as a block to form the paralagous two-gene clusters *Hmx3-Hmx2* and *Hmx1-SOHo* (Fig. 3E). Analysis of the broader chromosomal regions in which these genes lie has shown this occurred as part of the two rounds of genome duplication (2R) that occurred in early vertebrate evolution^22^.

**Figure 3.**
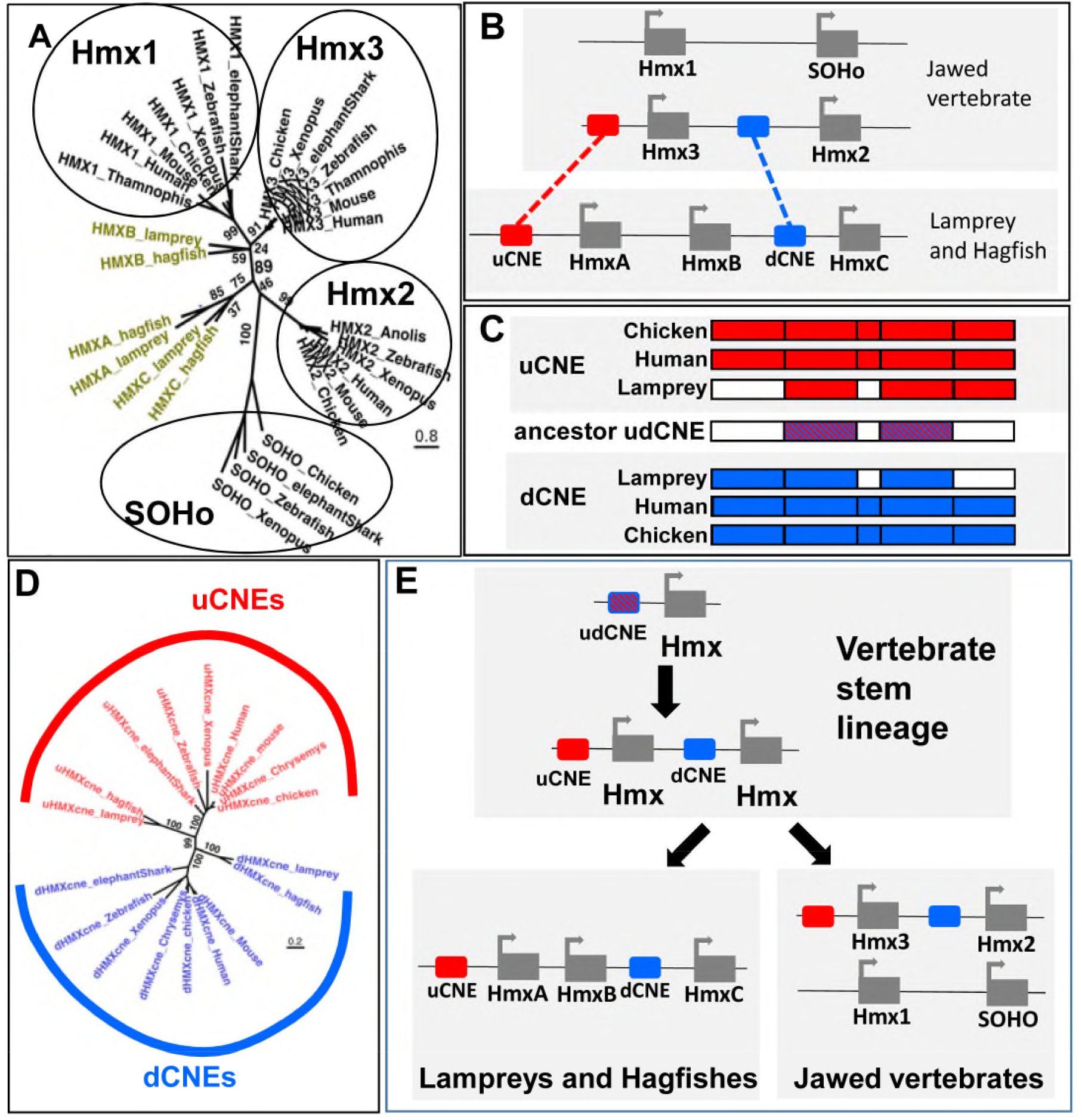
Vertebrate Hmx locus evolution and CNE identification. **A.** Phylogeny of vertebrate HMX proteins. Values indicate percentage bootstrap support. **B.** Comparative mapping of jawed and jawless vertebrate Hmx loci identifies two CNEs. **C.** Sequence similarity between uCNE and dCNE sequences in lamprey, human and chicken show they derived from an ancestral pre-duplication CNE. Coloured boxes indicate regions of sequence similarity amongst uCNE sequences (red) dCNE sequences (blue) and between uCNE and dCNE sequences (dashed red and blue). Sequence alignments in Supplementary Figure 1. **D.** uCNE and dCNE sequences form monophyletic groups in molecular phylogenetic analysis. **E.** A model for the evolution of Hmx loci. The current arrangements in vertebrates evolved from a single ancestral cluster with both uCNE and dCNE elements present, which itself evolved from a one-gene state with a single ancestral udCNE.

Comparative genomics across all jawed vertebrates revealed two CNEs, both associated with the *Hmx3-Hmx2* locus. One (uCNE) lies 5’ of *Hmx3*, the other (dCNE) lies between *Hmx3* and *Hmx2*, 5’ of *Hmx2* (Fig. 3B). Both are over 1kb long (Figure S1), exceptionally large for ancient vertebrate CNEs^23,24^. Unusually, they are also homologous to each other, sharing a core of around 500bp of highly conserved sequence between paralogous CNEs (Fig. 3C. S1). We searched for these CNEs in lamprey and hagfish, identifying one 5’ of *HmxA*, and a second between *HmxB* and *HmxC* (Fig. 3B; Fig. S1). Molecular phylogenetic analysis confirmed their orthology to jawed vertebrate uCNE and dCNE respectively (Fig. 3D). These data show the uCNE and dCNE originally evolved by tandem duplication of one ancestral CNE (udCNE: Fig. 3C, E). Parsimoniously this happened at the same time as the ancestral Hmx gene was duplicated, predating the separation of jawless and jawed vertebrate lineages over 500 million years ago^25^.

To test the functions of uCNE and dCNE we generated lamprey embryos transgenic for reporter constructs (Fig. 4A)^26^. Both lamprey CNEs drove reporter expression in the CNS in a pattern similar to endogenous Hmx gene expression (Fig. 4B-G). Confocal imaging showed uCNE was also able to drive reporter expression into CSG derivatives including the facial, glossopharyngeal and vagus nerves, as well as into some structures which do not derive from CSG or express Hmx (Fig. 4F-G’). These data confirm that the CNEs are regulatory elements which capture aspects of the spatial expression of lamprey Hmx genes. Since they evolved by duplication and divergence from udCNE, we infer this ancestral element would also have had the capacity to drive gene expression into CNS and CSG (Fig. 4H).

**Figure 4.**
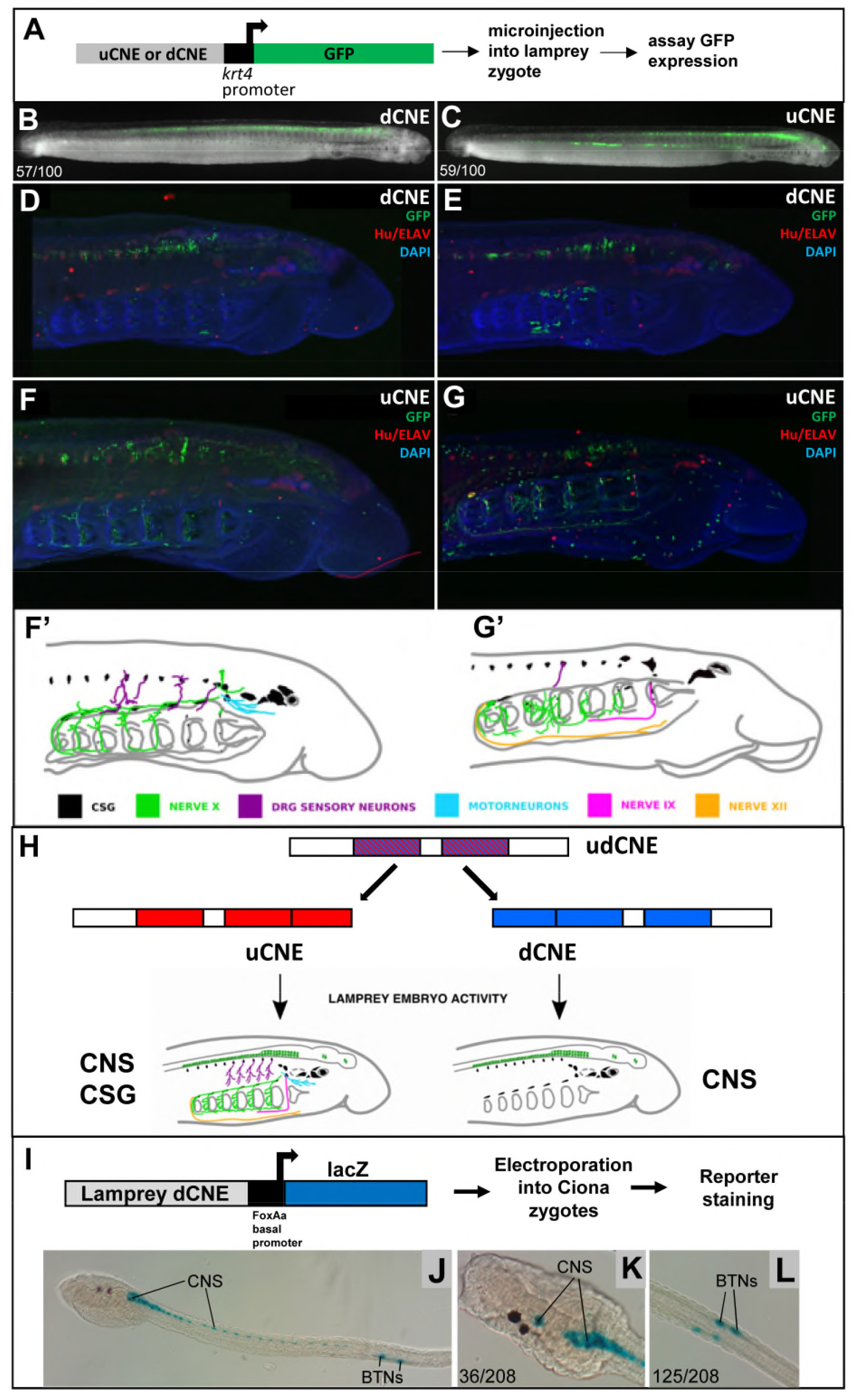
Lamprey Hmx CNE activity in transgenic lamprey and *Ciona* embryos. **A.** Experimental strategy for detecting reporter activity in lamprey embryos. **B, C.** Representative embryos showing *dCNE::GFP* and *uCNE::GFP* activity in the CNS. Numbers show the number of times CNS expression was seen out of the number of embryos screened. Analysis of vector-only controls is shown in Supplementary Figure S3. **D, E.** Confocal reconstructions of lamprey embryos transgenic for *dCNE::GFP*, showing activity (green) in forebrain, midbrain, hindbrain and spinal cord. **F, G.** Confocal reconstructions of lamprey embryos transgenic for *uCNE::GFP*, showing activity in the CNS, and in peripheral nerves. In D-G ganglia and other neurons are stained red using an antibody to Hu/ELAV. **F’, G’** Schematic tracing of embryos shown in F and G respectively, with nerve CNE reporter activity traced and colour coded. **H.** Summary of lamprey Hmx CNE activity. **I-L.** I shows a schematic of the experimental method used to examine lamprey CNE activity in *Ciona.* Below are transgenic larvae stained for lamprey dCNE reporter activity. Numbers indicate the number of times expression in the cells identified was seen, out of total surviving larvae. Full embryo counts are in Supplementary Figure S6.

The deduced ancestral vertebrate regulation of Hmx has similarity to the regulation of Hmx in living *Ciona*, with both including a large, proximal CNE driving expression into both PNS and CNS. This raises the possibility that regulation in the two lineages is homologous, in turn predicting BTNs and CSG will share the regulatory environment needed to activate Hmx expression. While the *Ciona* and vertebrate regulatory elements do not show sequence conservation, when we tested the activity of lamprey CNEs in transgenic *Ciona* (Fig. 4H) we found dCNE was able to recapitulate *Ciona Hmx* expression in BTNs and CNS (Fig. 4I-L). BTN expression was specific and robust (Fig. 4J, L). CNS expression extended along the majority of the neural tube (Fig. 4J), encompassing cells in the anterior *Ciona* CNS that express *Hmx*, but resembling the more extensive CNS expression of Hmx genes and reporters in amphioxus and vertebrates (compare Fig. 4I-L with Fig. 4B, C and Fig. 1I-M).

It has been previously suggested that BTNs are homologs of the neural crest^3^, and this has been elaborated into a gene regulatory model in which the *Ciona* neural plate border divides under an anterior-posterior (AP) patterning network into a posterior ‘proto-neural crest’ domain and anterior ‘proto-placode’ domain^2^. However, Hmx constitutively marks vertebrate placodes and CSG, and not neural crest: while Hmx expression in DRG and ciliary ganglia (both neural crest derivatives) has been reported in the mouse ^7,27,28^, this is not seen in other vertebrates including *Xenopus*, zebrafish, medaka and lamprey^6,8,9,13^ (Supplementary Figure S4). Our expression data point to BTNs as homologous to placodal CSG neurons and not to neural crest. This is reinforced by finding that a vertebrate Hmx CNE robustly drives expression of a reporter in *Ciona* BTNs and Lamprey CSG neurons, implying the two cell types share similar *trans*-regulatory environments. Furthermore, this matches the embryonic origin of both BTNs and placodes as lateral to the neural plate. *Ciona* also produces Hmx-negative PNS cells from the anterior neural plate border. These cells express markers of vertebrate chemosensory and GnRH neurons^2^, which in vertebrates develop from the olfactory placode. Our data are hence in keeping with proposals that the ancestral neural plate border had two domains yielding PNS cells^2,29^. Both, however, are homologous to placode-derived CSG, and not neural crest.

We also note evolution of Hmx in tunicates and vertebrates parallels derived aspects of neural evolution and includes highly unusual features. Tunicates are secondarily-reduced^30^, and we see direct evidence for how this evolved in the broad activity of vertebrate dCNE in the *Ciona* CNS. This shows the gene regulatory environment necessary for broad ancestral Hmx expression has persisted in *Ciona*, but that *Ciona Hmx* expression has become restricted to the anterior CNS by changes in its *cis*-regulation. Following divergence of the tunicate and vertebrate lineages, Hmx evolved by tandem duplication of gene and CNE in the stem vertebrate. That both CNEs have been subsequently conserved and maintained in tandem over the remainder of vertebrate evolution appears to be unique. These Hmx CNEs are also unusually large, with other identified jawed vertebrate CNEs conserved with lamprey typically much smaller ^24^. While the functional implications of these unusual features are unknown, such conservation speaks to extreme evolutionary constraint. We speculate this reflects a requirement for robustness in interacting with the ancient *trans* regulatory environment that evolved in the stem lineage of the tunicates and vertebrates, perhaps reflecting the instructive role of Hmx in directing the development of an ancient cell type involved in environmental sensing.

## Materials and Methods

### Hmx gene identification, cloning and sequence analyses

We searched multiple sources of lamprey and hagfish sequence data for potential Hmx genes. For lamprey this constituted *Lampetra planeri* transcriptome data ^1^, genome assemblies for *Petromyzon marinus*^2,3^, a *P. marinus* transcriptome assembly built in-house from Illumina GAII data available on SRA, the *Lethenteron camtschaticum* genome assembly ^4^, and an L. *camtschaticum* transcriptome assembly kindly provided by Juan Pascual-Anaya. For hagfish we searched the *Eptatretus burgeri* genome Eburgeri_3.2 genome assembly. In each dataset we identified the three genes as described in the main manuscript. In *P. marinus* (genome version Pmar_germline 1.0/petMar3) these are located on scaffold_00015 represented by gene models PMZ_0020818-RA, PMZ_0048148 and PMZ_0028877-RA. An additional *P. marinus* scaffold, scaffold_00813, also contained a gene model (PMZ_0038761-RA) with an Hmx type homeobox. However when we examined the sequence of this locus it was found to have >99.5% identity to part of the Hmx locus from scaffold_00015 (Figure S3, supplementary data). We concluded it is either a very recent duplication of sequence from scaffold_00015, or an artefact of the genome assembly process, and have not considered it further.

To identify CNEs we first compared jawed vertebrate loci using the Conserved Non-coDing Orthologous Regions (CONDOR) database ^5^. This identified a small number of elements surrounding the *Hmx3-Hmx2* locus that were conserved across jawed vertebrates. We extended this to lamprey and hagfish, using sequence similarity searches to search specifically for these conserved elements in these lineages. Alignments and molecular phylogenetic analyses were undertaken using MUSCLE ^6^ and RAxML ^7^ using the Maximum Likelihood method. 1000 bootstrap replicates were used to assess node confidence.

Lamprey *Hmx* genes were cloned from *L. planeri* cDNA, and uCNE and dCNE sequences from *L. planeri* genomic DNA, using the primers shown in the Table 1 below. The *Ciona* intestinalis *Hmx/Nkx5* locus was already annotated ^8^, though the gene model was incomplete. Since no ESTs mapped to this gene, no clones were available in arrayed plasmid libraries. We hence first cloned a fragment of the gene using the primers shown in the table below, and used this for in situ hybridisation. We then used homology to *Ciona savignyi*, coupled with RNAseq data mapped on ANISEED ^9^, to identify the full open reading frame. This was amplified, using the primers shown below, for the overexpression experiment. *Branchiostoma lanceolatum Hmx* was identified by searching the genome ^10^. The in situ probe was cloned by PCR from 24-36 hours post fertilisation larvae cDNA.

**Table 1:**
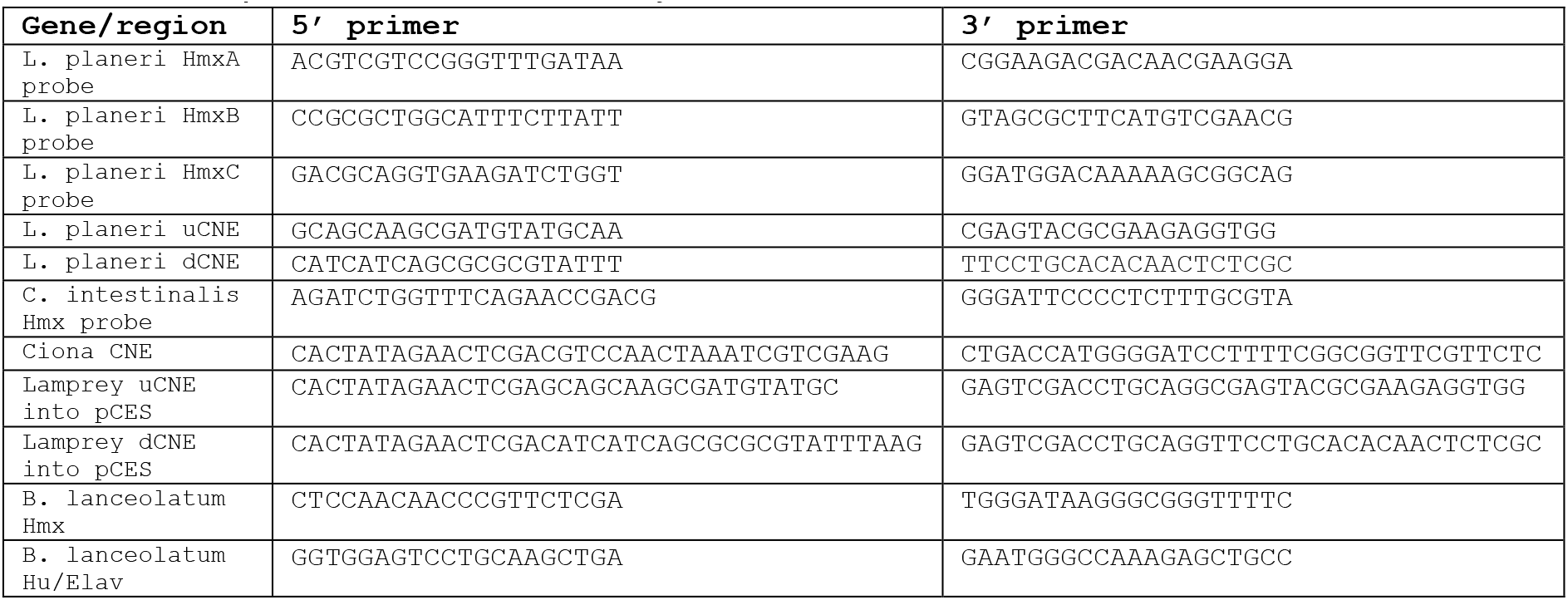
PCR primers used in this study

All clones were verified by sequencing, and new cloned sequences have been deposited in Genbank accessions MN264670-MN264672.

### Embryos and In situ hybridisation

Naturally spawned *Lampetra planeri* embryos were collected from a shallow stream in the New Forest, UK, under a Permission granted by Forestry England. They were cultured in filtered river water at 16°C and processed for in situ hybridisation as previously described ^11^. Adult *Ciona intestinalis* were collected from Northney Marina, UK, and maintained in a circulating sea water aquarium at 14°C under constant light. Gametes were liberated by dissection, fertilised in vitro and embryos allowed to grow to the desired stage before fixation and storage. Methods for fixation, storage and in situ hybridisation were as previously described ^12^. Adult *Branchiostoma lanceolatum* were collected near Banyuls-sur-Mer, France and spawning was induced by heat stimulation ^13,14^. Embryos were grown for 36hr at 19°C in natural sea water. Fixation was performed for 2hr on ice in 4% PFA in MOPS buffer containing 0.1M MOPS, 1mM EGTA, 2mM MgSO_4_ and 500mM NaCl. In situ hybridization was performed as previously described ^15^.

### Lamprey transgenics, imaging and controls

Lamprey uCNE and dCNE sequences from *L. planeri* were amplified by PCR (primers in Table 1) and cloned into the HLC vector with a zebrafish *krt4* minimal promoter ^16^. Lamprey transient transgenesis was performed in *P. marinus* embryos as previously described ^16,17^. Briefly, injection mixes consisting of 20ng μl^-1^ reporter plasmid, 1x CutSmart buffer (NEB), and 0.5U μl^-1^ I-SceI enzyme (NEB) in water were incubated at 37°C for 30 minutes and then micro-injected at a volume of approximately 2nl per embryo into lamprey embryos at the one-cell stage. Embryos were then raised and screened for GFP reporter expression using a Zeiss SteREO Discovery V12 microscope. Transient transgenic reporter assays may generate mosaicism in reporter expression patterns, with variation in levels and domains between embryos. 100 embryos were screened for each construct at two stages (25 and 27)

Representative GFP-expressing embryos were first imaged live to record GFP fluorescence, using a Zeiss SteREO Discovery V12 microscope and a Zeiss Axiocam MRm camera with AxioVision Rel 4.6 software. Embryos were then fixed in 4% paraformaldehyde and stained with a Chicken Polyclonal Anti-GFP antibody (Abcam AB13970),and a Mouse HuC/HuD Monoclonal Antibody (Invitrogen 16A11). These were visualised with Goat Anti-Chicken IgY H&L Alexa Fluor® 488 (Abcam AB150169) and anti-Mouse alexa594 (Abcam AB150116). Before imaging, embryos were counterstained with DAPI. Embryos were viewed on an Olympus FV1000 Confocal microscope. Reconstructions and analysis were carried out using FIJI-imageJ v.1.52g ^18^. Z stacks and 3D projects of confocal data were built using maximum intensity projection.

Confocal microscopy was able to reveal GFP expressing cells not possible to image in live embryos. Since lamprey transgenesis is a relatively new technique and previous studies have not assessed levels of background at this resolution, we analysed reporter activity in embryos injected with the plasmid vector HLC (Figure S2), focusing on ganglia and CNS expression that might overlap with endogenous *Hmx* staining and confound interpretation. This revealed GFP expression in skin cells, head muscle and branchiomeric muscle. We also saw occasional expression in CNS and ganglia. CNS expression was clearly distinct from that observed with *Hmx* enhancers and did not overlap with *Hmx* gene expression. Ganglia expression was infrequent (22% of embryos analysed: Figure S2, supplementary data) and did not label the same cells as seen with Hmx uCNE.

### *Ciona* transgenics and sequence analysis

The plasmid vector containing the *epiB* promoter driving *GFP (epiB::GFP)* was kindly provided by Bob Zeller ^19^. The full *Hmx* open reading frame was amplified by PCR and cloned downstream of the *epiB* promoter, replacing GFP and creating *epiB::Hmx. Ciona intestinalis* Hmx was amplified in two sections, a 5’ region using the primers TAAAATAGTAAAATGGTACCTATGACGTCACTGTGCCAATTG and TTCCCCTTCTGACGTAGGGA, and a 3’ section using the primers TCCCTACGTCAGAAGGGGAAG and ACCGGCGCTCAGCTGGAATTATGATTGTCTCACACCACGGAA. This resulted in two fragments: each had a homologous arm overlapping the other fragment and a homologous arm overlapping one end of the vector digested with KpnI and EcoRI. The Cold Fusion system (System Biosciences) was used to insert these into the vector via recombination, fusing the 5’ end of the resulting full *ciHMX* ORF with the 3’ end of the *epiB* promoter. Integrity of the resulting construct was confirmed with sequencing.

Constructs were electroporated into C*. intestinalis* zygotes as previously described ^20^. We first confirmed that these constructs drove their respective transgenes into the epidermis as expected, using GFP live imaging and *Hmx* in situ hybridisation respectively. We then electroporated parallel batches with either *epiB::GFP* only (control) or *epiB::GFP* and *epiB::Hmx (Hmx* overexpression) constructs. Embryos were grown to the tailbud stage when GFP was visible, identifying transgenic embryos which were then manually selected and processed for RNA extraction. Three full biological replicates were performed on embryo batches derived from different fertilisations, with each biological replicate combining RNA from at least 50 individual embryos. All six RNA samples were sequenced by Illumina HiSeq4000 following polyA selection, yielding approximately 28 million paired end 75bp reads per sample.

For differential gene expression analysis, fastQC was used to assess sequencing quality, Trimmomatic ^21^ to trim off adapters and Sickle ^22^ to trim low quality reads. Remaining reads were then mapped to the *Ciona robusta* genome (KH2012) using STAR ^23^. Differential expression analysis was carried out using the DESeq2 R package ^24^, using an adjusted p value threshold of 0.01. Finally, a minimum FPKM threshold of 2 was also applied to exclude very lowly expressed transcripts. This yielded a list of genes significantly (adjusted p<0.01) up or downregulated in the *Hmx* overexpression treatment compared to the control. Gene lists deriving from *C. intestinalis* single cell sequencing were extracted from the supplementary files of published literature ^25^, and cross-correlated with the up and down regulated gene lists to provide the annotation of data shown in Figure 2B.

To test CNE activity in *Ciona*, we cloned the 2kb 5’ to *C. intestinalis Hmx*, lamprey uCNE or lamprey dCNE into the reporter vector pCES ^26^ using the primers shown in Table 1. Constructs were electroporated into *C. intestinalis* zygotes as above, and embryos stained for β-galactosidase activity as described ^27^.

